# Neuro-metabolic pathways of high-protein meal reducing food craving

**DOI:** 10.64898/2026.01.06.697861

**Authors:** Min Pu, Rémi Janet, Sergio Oroz Artigas, Anja Ulrich, Jeremy Tardu, Peter N.C. Mohr, Britta Wilms, Berthold Koletzko, Sebastian M. Meyhoefer, Soyoung Q Park

## Abstract

Across species, high-protein foods reduce appetite throughout the day; however, their underlying neuro-metabolic mechanisms remain unknown. Here, we investigated whether a protein-rich diet modulates plasma tyrosine dynamics, a dopamine precursor, thereby altering brain activity in dopaminergic regions and reducing food craving in humans. In this highly controlled within-subject crossover study (preregistered: https://osf.io/du6h5), 30 healthy participants (age, 23.63 ± 3.23 years) received either a high- or low-protein breakfast. Three and a half hours after breakfast, participants viewed high-or low-calorie food images while undergoing magnetic resonance imaging (MRI), and their subjective food craving was assessed using the validated Food Craving Questionnaire. Plasma tyrosine levels were continuously monitored during the experimental procedure. Our results show that a high-protein breakfast significantly reduced subjective food craving compared with a low-protein breakfast. Notably, we found that a high-protein breakfast significantly enhanced plasma tyrosine levels, which were associated with lower food craving four hours after meal intake. Importantly, we observed that greater midbrain activity related to viewing food stimuli following the high-protein breakfast was associated with lower food craving. Together, these results provide strong evidence for a systemic body-brain interplay in regulating food craving throughout the day in humans.

**Graphic Abstract:**

- High-protein breakfast significantly reduces food craving over 4 hours compared to low-protein breakfast.
- High-protein breakfast significantly elevates plasma tyrosine levels, which were associated with lower food craving.
- High-protein breakfast increases midbrain activity significantly while viewing food images, predicting lower food craving.

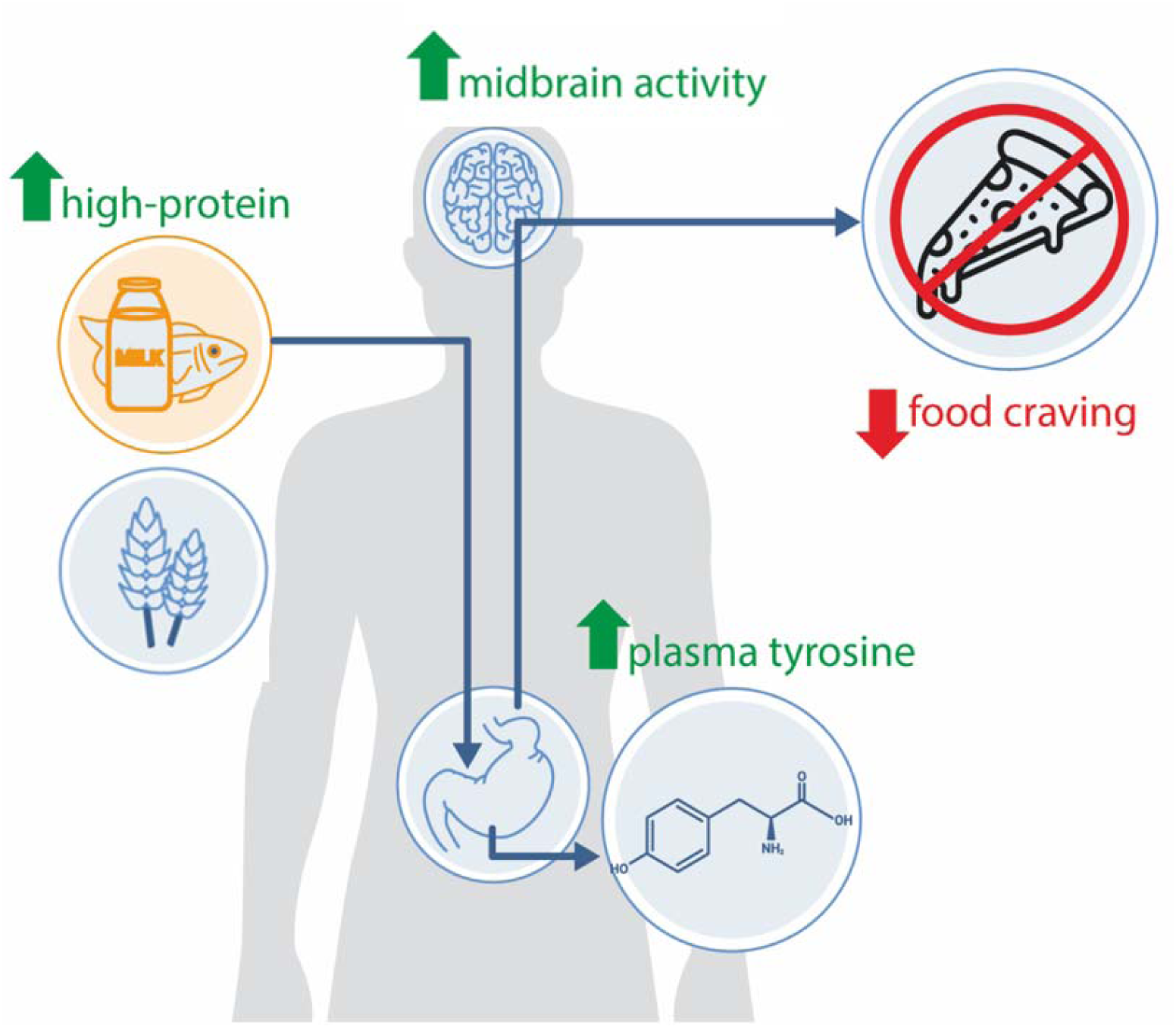

## INTRODUCTION

Compared with carbohydrates and fats, protein is considered the most satiating macronutrient and has a greater thermogenic effect ^1,2^. Previous research consistently shows that high-protein intake (20-30% of total energy intake) inhibits appetite by increasing satiety ^3^, decreasing the desire to eat, and reducing subsequent food intake in both humans ^4,5^ and animals ^6^. Importantly, the satiety effects of consuming a high-protein meal persist throughout day ^7^. For example, primates regulate their total food intake based on the protein content of their diet, allowing fat and carbohydrate intake to fluctuate ^8^. Specifically, when eating an imbalanced diet, such as one low in protein but high in fat and carbohydrates, primates show dysregulated intake of fats and carbohydrates, leading to energy excess or deficit. These studies suggest that high-protein intake helps maintain lean body mass and increases energy expenditure, whereas lower protein intake leads to compensatory food consumption.

Although the satiety effect of protein intake is well-documented, its underlying neural-metabolic mechanism remains unknown. Exploring this pathway is essential for understanding how protein impacts human food intake and has important implications for obesity research. One potential explanation for protein-induced satiety is the aminostatic hypothesis, which suggests that increased plasma amino acid levels following protein consumption enhance satiety ^9^. The macronutrient composition of foods alters the levels of large neutral amino acids (LNAAs) in the blood. Specifically, protein intake increases plasma tyrosine levels ^10^, the precursor of the neurotransmitter dopamine ^11,12^. Extensive evidence demonstrates that changes in peripheral tyrosine levels influence central dopamine function across species. In humans, positron emission tomography (PET) studies have shown that acute dietary tyrosine depletion via an oral amino acid mixture significantly decreases central dopamine release, as evidenced by increased [^11^C] raclopride binding in the ventral striatum ^13,14^. Importantly, these studies found that changes in dopamine release were significantly associated with peripheral tyrosine availability. Similarly, rats with reduced plasma tyrosine, induced by an amino acid mixture that omitted phenylalanine and tyrosine, showed significantly reduced dopamine transmission ^15^. In contrast, oral tyrosine supplementation facilitated dopamine metabolite availability in the rat brain ^16^. These findings collectively suggest a close link between plasma tyrosine and central dopamine responses, suggesting that tyrosine manipulation may serve as a potential mechanism for modifying dopamine-dependent behaviors such as craving.

Food craving is a psychological state that impacts multiple cognitive processes involved in food-seeking behavior, including attentional bias ^17^ and reward processing ^18^, and motivational drive ^19^. Facilitated dopamine availability is associated with reward processing, thereby influencing a range of human behaviors, including food consumption, motivation, and reward-related behaviors ^20^. Notably, the brain’s dopamine synthesis system plays a key role in regulating craving-related behaviors, including cigarette smoking, drug addiction ^21^, and food craving ^22^. In line with this, regulating plasma tyrosine concentration has been shown to alter craving, presumably through dopaminergic mechanisms. For example, in smokers, acute tyrosine depletion attenuates dopamine release ^23^ and increases cigarette consumption ^24^. Given dopamine’s central role in craving regulation, dopamine modulation may be a promising mechanism underlying the craving reduction effects of high-protein meals.

Previous studies have shown that a high-protein meal regulates human social behavior ^25^, which is driven by changes in tyrosine availability. Building on these findings, this present study aimed to examine how acute high-protein intake influences subjective food craving, as assessed by the validated Food Craving Questionnaire ^26^. We expected that high-protein meal-induced changes in food craving would be accompanied by changes in brain activity. Our goal was to understand the neuro-metabolic mechanisms through which a high-protein meal reduces food craving over time. We hypothesized that (1) a high-protein breakfast would decrease self-reported subjective food craving and increase plasma tyrosine dynamics across the day, and (2) plasma tyrosine would be negatively associated with subjective food craving after a high-protein breakfast. Since plasma tyrosine modulation is linked to the regulation of central dopamine levels, we hypothesized that (3) eating a high- vs. low-protein breakfast leads to greater brain activity in dopaminergic regions when viewing high-caloric food stimuli, associated with lower food craving. The hypotheses of this study were pre-registered on the Open Science Framework (https://osf.io/du6h5).

## RESULTS

To investigate the neuro-metabolic pathway underlying high-protein meal-induced changes in food craving, we conducted a randomized, counterbalanced, within-subjects study. We included 30 male participants in the analysis (age, mean ± SD, 23.63 ± 3.23 years; BMI, mean ± SD, 22.90 ± 1.80 kg/m^2^; see Methods for details). Participants visited the laboratory twice, approximately one week apart. Upon arrival, participants received either a high-protein breakfast (25% protein, 50% carbohydrate, and 25% fat) or a low-protein breakfast (10% protein, 80% carbohydrate, and 10% fat). Both meals included various food items, provided an equal energy content of 850 kcal, and were served at 08:45 h. To assess sustained dietary effects (high- and low-protein breakfast), we measured brain activity 3 hours after breakfast using functional magnetic resonance imaging (fMRI), while participants viewed high- and low-calorie visual food stimuli during a food choice task. Participants then completed a food craving questionnaire approximately 4.5 hours after breakfast **(Figure 1).** We assessed plasma tyrosine at four time points over five hours. Both sessions followed identical experimental procedures and differed only in breakfast composition (e.g., high- vs low-protein), presented in randomized order.

**Figure 1.**
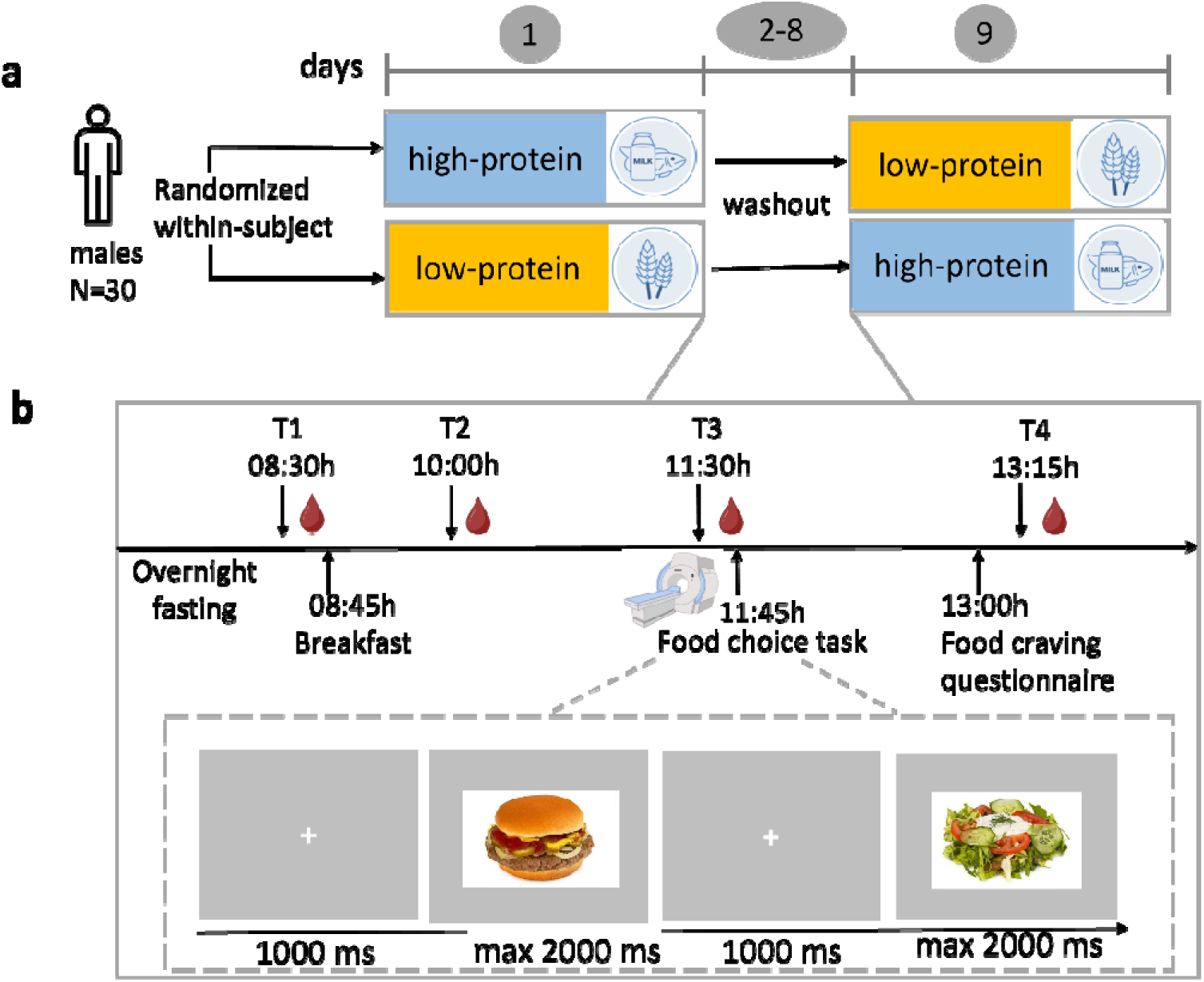
Experimental design and procedure. **a.** Experimental design. Randomized, counterbalanced, within-subject study. Participants visited the lab twice, about one week apart. They received either a high-protein breakfast (25% protein, 50% carbohydrate, and 25% fat) or a low-protein breakfast (10% protein, 80% carbohydrate, and 10% fat), each containing various food items totaling 850 kcal at 08:45 h. **b.** Experimental procedures. Participants consumed breakfast at 08:45 h. We collected blood samples at 08:30 h, 10:00 h, 11:30 h, and 13:15 h. Participants completed the food choice task at 11:45 h. In this task, participants viewed food images and decided whether they wanted to consume the items right away. Participants completed the food craving questionnaire at 13:00 h.

### Sustained impact ofa high-protein meal on food craving and metabolic dynamics

We first tested whether the high-protein breakfast reduced subjective food craving 4,5 hours after consumption using a paired t-test. As expected, participants reported significantly lower food craving after the high-protein breakfast compared with the low-protein breakfast (t (29) = 2.40, p = 0.02, Cohen’s d = 0.44) **(Figure 2a)**. Next, we tested whether the high-protein breakfast increased plasma tyrosine dynamics. A Wilcoxon test showed a significant effect of diet (high-and low-protein breakfasts), with higher tyrosine after the high-protein breakfast than after the low-protein breakfast, z = −2.48, p = 0.01. Next, we tested whether the high-protein meal altered tyrosine/LNAA at specific time points. A two-way repeated-measures ANOVA showed a significant interaction between diet and time points, F(2.11, 61.31) = 4.46, p = 0.014. Post-hoc analyses showed that compared to eating the low-protein breakfast, the high-protein breakfast induced significantly higher plasma tyrosine/LNAA levels at 11:30 (t(29) = −3.69, p (FDR) = 0.0009) and at 13:15 (t(29) = −3.07, p(FDR)= 0.0046; **Figure 2b**).

**Figure 2.**
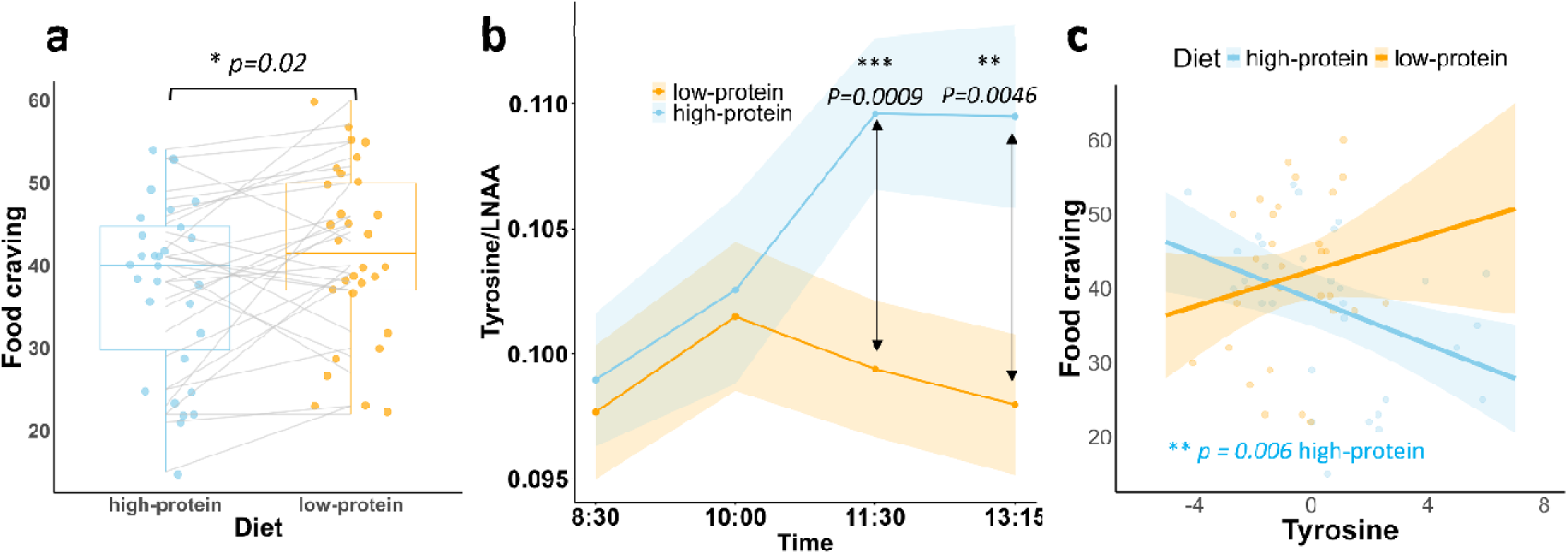
Impact of the high-protein breakfast on food craving and tyrosine. **a.** 4,5 hours after the high-protein breakfast, participants reported lower food craving. **b.** Plasma tyrosine/LNAA was significantly higher after the high-protein vs. low-protein breakfast and remained sustained at 13:15 h. **c.** Tyrosine dynamics were significantly related to (predicted) food craving after the high-protein breakfast. Tyrosine was quantified as the area under the curve of the tyrosine/LNAA ratio relative to the increase (See Methods for details). Tyrosine in Figure 2c was mean-centered. * p< 0.05, ** p<0.01, *** p< 0.001.

Next, we tested whether increases in tyrosine were associated with reduced craving following the high-protein breakfast. A linear mixed-effects model revealed a significant interaction between diet and tyrosine (β = −2.73, SE = 0.96, p = 0.007). Specifically, post-hoc analyses showed that after the high-protein breakfast, greater tyrosine was related to lower food craving (b = −23, SE= 8.03, p (FDR)= 0.006), whereas this link was absent following the low-protein breakfast (b = 18, SE= 14.24, p (FDR)= 0.21; **Figure 2c**). To rule out potential influences of ghrelin and insulin on food craving, we also ran an alternative model that included both variables as covariates. Our results showed that the interaction between diet and tyrosine remained significant (b = 2.91, SE = 1.08, p = 0.011). Additionally, paired t-tests revealed no significant difference in ghrelin between the two breakfast sessions (t = −0.46, p = 0.65, Cohen’s d = 0.09). Wilcoxon tests found no significant difference in insulin between breakfast sessions (z=0.93, p=0.37, Cohen’s d = −0.25). These results indicate that the craving reduction induced by a high-protein meal is driven by plasma tyrosine, independent of ghrelin and insulin.

### High-protein breakfast enhances midbrain activity, associated with lower food craving

Next, we investigated whether the neural response to palatable food after eating high- vs. low protein breakfast is related to craving reduction. To do this, we conducted whole-brain regression analyses in which we predicted the brain activity differences (high vs low calorie visual cues) using individual food craving differences between high vs low-protein breakfasts. We observed that a greater midbrain activation when reviewing palatable food pictures, after eating a high- vs. low-protein breakfast, was associated with a greater reduction in subjectively reported food craving (MNI coordinate [x=-4, y=-18, z=-16], p(Family-Wise Error-Small Volume Correction) < 0.05, t= 4.06 (**Figure 3**)). This indicates that protein-driven midbrain activation was directly related to craving modulation, as participants with greater midbrain upregulation also reported lower food craving when comparing high- vs. low-protein breakfast conditions (b = −6.84, t = −3.46, p = 0.0018). We did not observe significant brain activation in any other pre-registered dopaminergic regions (including the caudate, putamen, nucleus accumbens (NAcc), and orbitofrontal cortex (OFC). We also examined neural activation associated with viewing food stimuli (e.g., high-calorie > low-calorie) for each breakfast condition, which is shown in the supplementary materials.

**Figure 3.**
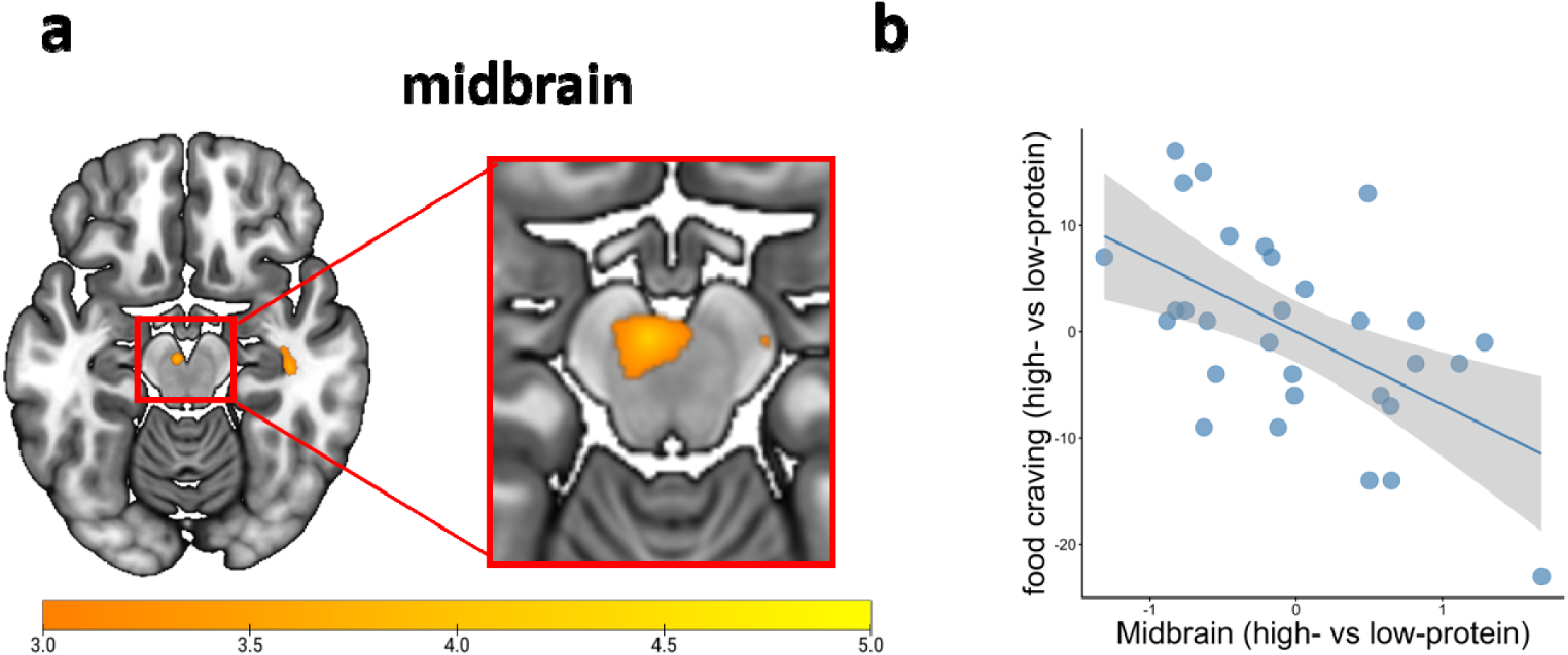
Neural differences between high- and low-protein breakfasts predict differences in subjective food craving. **a.** Whole-brain activity using small-volume correction (SVC) in the Midbrain (MNI coordinates [x=-4, y=-18, z=-16], cluster= 22, p(FWE-SVC) < 0.05). b. The correlation between the difference in parameter estimates of two breakfast conditions (e. g., high-protein (high >low calorie) - low-protein (high > low calorie)) and the difference in subjective craving rating. We also tested in other pre-selected dopaminergic brain regions, including caudate, putamen, nucleus accumbens (NAcc), and orbitofrontal cortex (OFC), which were not significantly modulated.

## Discussion

The present study investigated how a high-protein breakfast impacts food craving and its underlying metabolic and neural mechanisms. We found that the high-protein breakfast significantly reduced food cravings and increased peripheral tyrosine levels, compared with the low-protein breakfast. Notably, plasma tyrosine dynamics were negatively associated with food craving, depending on the protein-to-carbohydrate ratio of breakfast. Furthermore, the upregulation of midbrain activity after eating a high-protein breakfast led to reduced food craving. Together, these findings highlight essential metabolic and neural mechanisms that may explain how a protein-rich meal reduces reward-driven eating behavior.

The micronutrient composition of a meal impacts energy intake, metabolic function, and long-term body composition. Adequate protein intake is required for maintaining biological functions and health, whereas protein deficiency can lead to adverse health outcomes. Consistent with our hypothesis, we observed that the high-protein breakfast was associated with significantly lower food craving in healthy participants. Our findings align with previous research showing that high-protein diets increase satiety and reduce appetite ^27^. Even short-term high-protein diets have been found to reduce energy intake and body weight, likely through increased satiety and suppressed food intake in both animal ^6^ and human studies ^28,29^. However, while the mechanism of protein intake-related appetite regulation is well established ^30^, the present study was not designed to directly examine protein-specific appetite regulation. Instead, it complements this framework by examining transient subjective food craving and the underlying neuro-metabolic mechanisms following an acute protein intake.

As predicted, tyrosine levels were significantly higher after the high-protein breakfast than after the low-protein breakfast. Notably, greater peripheral tyrosine levels were associated with lower food craving following the high-protein breakfast, whereas this effect was absent after the low-protein breakfast. This effect remained significant after controlling for ghrelin and insulin levels. Moreover, in our study, the high-protein breakfast did not significantly increase plasma ghrelin levels, and ghrelin was not related to food craving in either breakfast session. Our findings challenge the traditional view that ghrelin, the “hunger hormone”, is the primary regulator of protein-induced satiety. Previous research has suggested that one of the main mechanisms of protein-induced satiety involves suppression of orexigenic hormones such as ghrelin ^31^. High-protein meals can suppress ghrelin release at the peripheral levels ^32,33^, which has been hypothesized to decrease appetite and subsequent food intake ^34,35^. However, these studies did not directly correlate ghrelin levels with food intake or subjective appetite, making conclusions inconclusive. Moreover, some studies have observed that high-protein induced appetite reduction is not associated with ghrelin homeostasis, particularly in obesity ^36^. In that study, participants were administered a weight-loss intervention, receiving either a high-protein, low-fat diet or a standard-protein, high-fat diet. The high-protein intervention decreased appetite independently of weight loss, whereas plasma ghrelin levels remained unchanged. Similarly, another study showed that a high-protein diet did not significantly alter fasting or postprandial ghrelin levels, despite reducing hunger ratings, and ghrelin was not correlated with hunger ratings ^37^. While these studies showed mixed results, our study provides novel evidence suggesting that plasma tyrosine dynamics represent an alternative pathway through which protein intake influences food craving.

A potential pathway linking high-protein induced craving reduction involves tyrosine-modulated dopaminergic signaling. Food macronutrient composition can alter metabolic signals. Specifically, digestion of high-protein meals breaks down proteins into amino acids such as tyrosine, a precursor of dopamine ^38^. In contrast, meals with a high-carb/protein ratio modify plasma tryptophan levels, the precursor of serotonin ^10^. Peripheral tyrosine availability is supposed to regulate dopamine synthesis in the brain ^39^. One mechanism underlying this relationship involves competition among large neutral amino acids (LNAAs) for transport across the blood-brain barrier, as they share the same saturable transporter. Increasing plasma tyrosine levels reduces the uptake of other LNAAs into the central nervous system (CNS), thereby increasing brain dopamine synthesis. In line with this mechanism, previous studies have indicated that macronutrient composition (e.g., high-protein), rather than protein intake alone, regulates social behaviors and decision-making ^25,40^. Our findings further suggest that high-protein meals can alter food cravings. In addition to this competitive transport mechanism, other pathways involving vagal signaling and gut-brain interactions may also link peripheral tyrosine to central brain dopamine function. The relationship between tyrosine and central dopamine function is well established. Acute phenylalanine/tyrosine depletion has been shown to alter reward evaluation ^41^ and to reduce food-related reward activity in the striatum ^42^ and nucleus accumbens^43^. Conversely, tyrosine supplementation improved brain dopamine availability and impacted cognitive function and behaviors ^44^. Our findings indicate that transient differences in dopaminergic responsivity, as reflected by tyrosine fluctuations, may contribute to individual variability in subjective food craving.

In our study, greater change in midbrain activity during viewing food stimuli was associated with a greater reduction in subjective food craving, after eating a high- vs. low-protein breakfast. This aligns with previous animal and human literature showing that central dopaminergic brain areas play a critical role in regulating food reward ^45^. Animal studies suggest that enhanced DA activity in midbrain regions is related to reduced food-related behaviour ^46^. For instance, chemogenetic activation of dopamine neurons in the midbrain was associated with decreased meal size and shorter meals in rats ^47^. In contrast, dopamine deficit in the mesolimbic circuit was associated with greater caloric intake and compensatory eating behavior in rats ^48,49^. In humans, acute phenylalanine/tyrosine depletion decreased food-related activity in reward-associated brain regions ^50^. Other studies also revealed that postprandial neural responses in the dopamine-related regions to food stimuli were positively correlated with fullness and reduced motivation to consume energy-dense foods ^51^, reflecting satiety-related changes in reward processing. Likewise, dopaminergic reward pathways are also modulated by homeostatic signals related to energy balance ^52^. In line with these studies, increased midbrain activity in response to high-calorie food stimuli following a high-protein breakfast does not necessarily indicate increased desire and wanting for food. But instead, with sufficient nutrient intake, increased midbrain activity may reflect state-dependent updating of food reward or altered motivational processing, both of which are linked to lower subjective craving.

Moreover, while our study focused on the acute effects of a single high-protein meal, a previous study examined the chronic effects of a low-protein diet over 16 days and reported enhanced reward-related neural activation in response to savory food cues ^53^. These findings suggest that protein-related neural responses may dynamically shift across physiological states, from enhanced reward responsivity following long-term protein restriction to stabilized dopaminergic function after sufficient acute protein intake. Additionally, future research should manipulate savory food cues associated with protein content to examine how acute protein intake influences brain responses related to taste-category preference and protein-specific content ^53^.

This study primarily tested pre-registered hypotheses regarding tyrosine as a physiological marker of a high-protein meal. However, other metabolic parameters, such as fibroblast growth factor 21 (FGF21) and glucagon-like peptide 1 (GLP-1), may also be modulated by protein intake. Research has revealed that protein restriction is associated with increased levels of FGF21 ^54^, suggesting that FGF21 induced by a low-protein diet may contribute to heightened food craving. Additionally, a high-protein meal has been found to increase GLP-1 ^55^. Future research should investigate whether, and through which mechanisms, FGF21 and GLP-1 may regulate the relationship among protein intake, food craving, and underlying neural processes.

While our findings primarily reflect the acute effects of a single high-protein meal, they cannot be generalized to habitual long-term protein intake. Habitual protein intake may exert distinct or additional influences on amino acids and appetite regulation through longer-term endocrine and metabolic adaptation ^30^. For instance, previous studies found that sustained high-protein diets over two weeks significantly reduce overall caloric intake through homeostatic adaptations involving metabolic hormones such as leptin ^56^.

Taken together, our study provides the first evidence of a neural metabolic mechanism by which a high-protein meal reduces food craving. Specifically, a high-protein breakfast significantly increased plasma tyrosine dynamics, which were associated with reduced food craving. At the neural level, the greater change in midbrain activity after eating a high-protein breakfast was associated with greater craving reduction. These findings have critical implications for understanding how nutrition modulates food intake and craving, and for developing dietary strategies aimed at prevention and weight management in obesity.

## METHODS

### Participants

An a priori power analysis was conducted using G*Power 3.1 to determine the required sample size for a paired-samples t-test. The analysis indicated that a total sample of 24 participants would be required to detect a medium effect size (d = 0.65) on food craving, with 85% power at an α level of .05 (two-tailed). This study recruited thirty-eight healthy male participants. However, eight participants were excluded from the final analysis: three failed to complete both fMRI sessions, one had abnormal structural brain images, one had missing tyrosine data, one exhibited excessive head motion during the fMRI task, and two failed to follow task instructions. Thus, the final sample included 30 participants (age, mean ± SD, 23.63 ± 3.23 years; BMI, mean ± SD, 22.90 ± 1.80 kg/m^2^). Furthermore, because four additional participants had missing values for both ghrelin and insulin, we included 26 subjects in the exploratory analysis using models for ghrelin and insulin. All participants provided written informed consent in accordance with the Declaration of Helsinki. The local Medical Ethical Commission of the University of Lübeck approved the study.

### Experimental design

The study procedure is displayed in **Figure 1a**. This was a randomized, counterbalanced, within-subject design consisting of two sessions separated by 7 to 9 days. Participants received either a high-protein breakfast (50% carbohydrates, 25% fats, and 25% proteins) or a low-protein breakfast (80% carbohydrates, 10% fats, and 10% proteins) in each session, with both meals providing an equal 850 kcal. As previous studies have suggested that the protein-to-carbohydrate ratio enhances central tyrosine levels through competition with other large neutral amino acids ^10,25,40^, we varied both protein and carbohydrate in the diets.

#### Blood samples

Fasting blood samples were obtained at 08:30 h, and participants consumed breakfast at 08:45 h. Additional blood samples were drawn at 09:00 h, 09:15, 09:30 h, 10:00 h, 10:30 h, 11:30 h, and 13:15 h. All samples were centrifuged at 4 °C and stored at −80 °C until analysis. Twenty-two plasma amino acids and ghrelin were measured in samples collected at 08:30, 10:00, 11:30, and 13:15 h using the method described ^57^. This process determined all proteinogenic amino acids, citrulline, and ornithine through a combination of precipitation, derivatization, and chromatographic separation ^40^. Insulin was measured at all eight time points using immunoassays (Immulite 2000, Siemens Healthcare Diagnostics, Erlangen, Deutschland). Ghrelin was assessed by radioimmunoassay (RIQ Kit, EMD Millipore Corporation, St. Louis, Missouri, USA).

#### Food choice task

Participants completed a food choice task inside the MRI scanner at 11:45 h. Each trial began with a fixation cross displayed for 1000 ms, followed by a food image. Participants indicated whether they were willing to consume the displayed food item within a 2000 ms response window by pressing a button (“Yes” or “No”). “Yes” and “No” responses were counterbalanced across the left and right sides across trials. No responses or responses after 2000 ms were counted as missed trials. The task consisted of four runs of 40 trials each (160 trials in total). Each run consisted of 20 high-calorie and 20 low-calorie food images, presented in random order. Two sets of pictures (each including 20 high-calorie and 20 low-calorie food images) were presented twice across the four runs.

#### Assessment of food craving

At 13:00 h, participants completed the German version of the State Food Craving Questionnaire (FCQ-S)^26^. The 15-item state version of the FCQ-S assesses momentary food cravings across five dimensions: intense desire to eat (desire), anticipation of positive reinforcement from eating (positive reinforcement), anticipation of relief from negative states or feelings as a result of eating (relief), lack of control over eating (lack of control), and craving as a physiological state (hunger). A total score was calculated.

### MRI data acquisition and preprocessing

MRI data were collected using a 3T MRI system (Siemens Magnetom Skyra, Germany) with a 64-channel head-coil. High-resolution anatomical images were acquired using a T1-weighted 3D MPRAGE sequence [TR = 1900 ms, TE = 2.44 ms, TI = 900 ms, FOV = 256 mm, volxe size = 1 x 1 x 1 mm]. Functional images were acquired using a T2*-weighted Gradient echo-planar images (EPI) with 58 slices (voxel size = 3 × 3 × 3 mm^3^, field of view (FOV)= 192 mm, repetition time (TR)= 2000 ms, echo time (TE)= 30 ms echo time, no distance factor), aligned parallel to the anterior commissure-poster commissure line (AC-PC) in an interleaved manner. Preprocessing, subject-level modeling, and group-level analyses were conducted using Statistical Parametric Mapping (SPM12; Functional Imaging Laboratory, London, U.K.). Functional imaging data were slice timing corrected, spatially realigned, inspected for excessive head motion (scans with displacement ≥ 3.0 mm were excluded), and normalized to the standard Montreal Neurological Institute (MNI) EPI template ^58^ (voxel size = 3 × 3 × 3 mm^3^). Normalized data were then spatially smoothed with an 8 mm isometric full width at half-maximum (FWHM). Subject-level models included event onsets convolved with the hemodynamic response function and six motion parameters from realignment, and models were high-pass filtered at 128 s.

### Imaging data analysis

The general linear model (GLM) implemented in SPM 12 was used to analyze the FCR task. At the first level of analysis, separate GLMs were created for the high-protein and low-protein sessions. Firstly, to examine dietary effects during the food choice task, onsets were specified at stimulus presentation, with two regressors of interest: high-calorie (HC) and low-calorie (LC) food images. We focused on the contrast of high-calorie > low-calorie. Afterward, first-level contrast maps were entered into a second-level random-effects analysis using a paired t-test (i.e., high-calorie vs. low-calorie). To examine whether differences in neural responses to food stimuli following a high-protein breakfast were related to changes in food craving, we conducted a second-level whole-brain regression analysis. The dependent variable was the contrast image reflecting the difference in viewing food stimuli between the two breakfast conditions [high-protein (high vs low calorie) and low-protein (high vs low calorie)], and the independent variable was the difference in food craving between the high vs low protein conditions.

#### Whole brain and regions of interest analyses (ROI)

Regions of interest (ROIs) associated with dopaminergic functions were defined using midbrain ROIs, caudate, putamen, nucleus accumbens (NAcc), and orbitofrontal cortex ^59^. Significant brain activation maps were defined at the voxel level of pÖ<Ö0.005, uncorrected, with a minimum cluster size of 10 voxels, and clusters were restricted with a family-wise error (FWE) correction at a cluster-wise threshold of pÖ<Ö0.05. For the high-protein versus low-protein contrast, given the prior hypothesis, a small-volume correction (SVC) was applied to identify significant brain activations within predefined ROIs. Dopaminergic-related region ROIs were derived from previous studies, as preregistered. The ROIs for midbrain, caudate, NAcc, and putamen were defined as 8 mm spheres centered at the peak value (midbrain: −4 −14 −12, 4 −14 −12; caudate: 12 10 −8, −6 12 −4; putamen: −14 8 0) ^60–63^. Parameter estimates for each contrast were extracted from these significant clusters.

## STATISTICAL ANALYSIS

To investigate whether a high-protein breakfast impacted food craving and tyrosine levels, paired t-tests compared high-protein and low-protein sessions for food craving, tyrosine, insulin, and ghrelin. The Wilcoxon test was conducted when the data were non-normally distributed. A two-way repeated-measures ANOVA was conducted to examine the effect of diet (high vs. low-protein breakfast) and time (08:30, 10:00, 11:30, 13:15) on the tyrosine/LNAA ratio. We next conducted linear mixed-effects models to assess the fixed effects of diet (high-protein vs. low-protein) and tyrosine on food craving, including their interaction. Participant ID was modeled as a random effect to account for repeated measures within subjects. The “emmeans” package and simple slopes analysis were used to examine significant interaction effects. Where necessary, false discovery rate (FDR) correction was applied to control for multiple testing. For exploratory analysis, to control for the potential impact of ghrelin and insulin on food craving, a second linear mixed-effects model included diet, tyrosine, their interaction term, ghrelin, and insulin as covariates. To investigate whether neural signals induced by the high-protein breakfast were associated with behavioral changes, linear regression analyses were conducted on difference scores (high minus low-protein) for food craving and midbrain beta values.

### Tyrosine calculation

Tyrosine was calculated as the ratio of plasma tyrosine concentration to the sum of the concentrations of five other large neutral amino acids (LNAAs), serving as a proxy for brain tyrosine and dopamine levels ^10^. The ratio was calculated using the following equation.

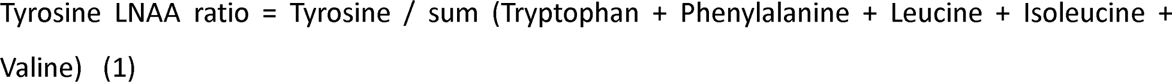

### Tyrosine area under curve (AUC) calculation

To quantify meal-related changes in tyrosine/LNAA ratio, trapezoidal area under the curve with respect to increase (AUC_I_) was calculated:

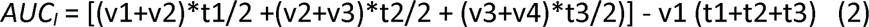

Where v is the tyrosine/LNAA ratio at a given time point t, and t1 to t3 are the intervals between measurements. Ghrelin AUC_I_ and insulin AUC_I_ were calculated in the same way. For simplicity, tyrosine/LNAA AUC_I_, ghrelin AUC_I_, and insulin AUC_I_ were referred to as tyrosine, ghrelin, and insulin, respectively, throughout this paper.

## Declaration of interests

The authors declare no competing interests.

## Acknowledgements

This study was funded by the Bundesministerium für Bildung und Forschung (BMBF; grant 01EE2301E) as part of the concept development of the German Center for Mental Health (DZPG), Deutsches Zentrum für Diabetesforschung (DZD grant 82DZD03D03, 82DZD03E5G) and the State of Brandenburg.

Data Availability Declaration

The data reported in this paper will be shared by the corresponding author upon reasonable request.

## AUTHOR CONTRIBUTIONS

**Min Pu:** Data curation, Formal analysis, Methodology, Visualization, Writing – original draft, Writing – review & editing. **Rémi Janet:** Formal analysis, Methodology, Writing – review & editing. **Sergio Oroz Artigas:** Investigation, Data curation,Writing – review & editing. Writing – review & editing. **Anja Ulrich:** Investigation, Data curation, Writing – review & editing. **Jeremy Tardu:** Investigation, Data curation,Writing – review & editing. **Peter N.C. Mohr:** Methodology, Writing – review & editing. **Britta Wilms:** Methodology, Software, Writing – review & editing. **Berthold Koletzko:** Conceptualization, Methodology, Writing – review & editing. **Sebastian M. Schmid:** Conceptualization, Funding acquisition, Writing – review & editing. **Soyoung Q. Park:** Conceptualization, Methodology, Supervision, Funding acquisition, Writing – original draft, Writing – review & editing.

## Notes

### Competing Interest Statement

The authors have declared no competing interest.

### Summary of Updates

Figure 3 has been updated, and euclideance distance analysis has been updated.

