## supplemental material for "Neuro-metabolic pathways of high-protein meal reducing food craving"

**Supplementary materials**

**The neural activity of food cue reactivity after eating the high-protein breakfast**

We examined neural activation associated with food cue reactivity (e.g., high-calorie > low-calorie) for each breakfast condition. After eating the high-protein breakfast **(Figure S1a),** participants showed significantly stronger brain activations in the precuneus (MNI[22 -50 44], p (FWE) < 0.001, t=5.26), inferior temporal gyrus (MNI[48 -58 -6], p (FWE) < 0.001, t=5.24), postcentral gyrus (MNI[-32 -40 48], p (FWE) < 0.001, t=4.45), and mediodorsal medial magcellular (MNI[2 -8 2], p (FWE) =0.028, t=5.24). After eating the low-protein breakfast **(Figure S1b),** participants showed significantly stronger brain activations in the fusiform gyrus (MNI[-44 -62 -14], p (FWE) < 0.001, t=4.97) and superior temporal gyrus (MNI[-58 -6 4], p (FWE) =0.022, t=4.74). However, the group comparison between high-protein and low-protein diets did not reveal any significant brain activations.

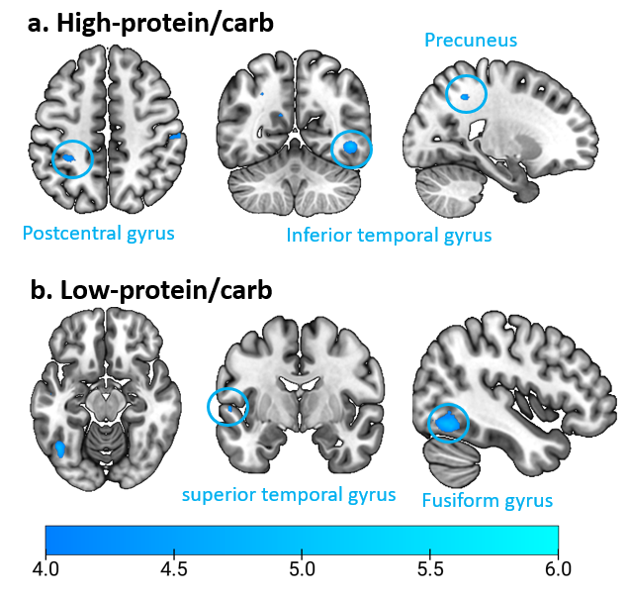

**Figure S1. Neural response to high-calorie food stimuli for both breakfast conditions. a.** Whole brain activity of the contrast high versus low- calorie in the high-protein condition. b. Whole brain activity of the contrast high versus low- calorie in the low-protein condition. Significant brain activation maps were defined at the voxel level of p < 0.005, uncorrected, with a minimum cluster size of 10 voxels, and clusters were restricted with a family-wise error (FWE) correction at cluster-wise threshold p < 0.05.
